# Identification and characterization of two transmembrane proteins required for virulence of *Ustilago maydis*

**DOI:** 10.1101/2021.03.09.434588

**Authors:** Paul Weiland, Florian Altegoer

## Abstract

Smut fungi comprise a large group of biotrophic phytopathogens infecting important crops such as wheat and corn. Through the secretion of effector proteins, the fungus actively suppresses plant immune reactions and modulates its host’s metabolism. Consequently, how soluble effector proteins contribute to virulence is already characterized in a range of phytopathogens. However, membrane-associated virulence factors have been much less studied to date. Here, we investigated six transmembrane (TM) proteins that show elevated gene expression during biotrophic development of the maize pathogen *Ustilago maydis*. We show that two of the six proteins, named Vmp1 and Vmp2 (**v**irulence-associated **m**embrane **p**rotein), are essential for the full virulence of *U. maydis*. The deletion of the corresponding genes lead to a substantial attenuation in the virulence of *U. maydis*. Furthermore, both are conserved in various related smuts and contain no domains of known function. Our biochemical analysis clearly shows that Vmp1 and Vmp2 are membrane-associated proteins, potentially localizing to the *U. maydis* plasma membrane. Mass photometry and light scattering suggest that Vmp1 mainly occurs as a monomer, while Vmp2 is dimeric. Notably, the large and partially unstructured C-terminal domain of Vmp2 is crucial for virulence while not contributing to dimerization. Taken together, we here provide an initial characterization of two membrane proteins as virulence factors of *U. maydis*.

## 1 Introduction

An increasing number of infectious diseases are threatening agricultural and natural systems. This development results in large crop losses, with up to 20 % of maize harvest loss caused by fungal pathogens such as *Ustilago maydis* (Fisher et al. 2012). Despite the high number of fungal species infecting plants, only a few fungal plant pathogen systems allow the physiological, molecular, and biochemical investigation of both host and parasite (Dean et al. 2012; Giraldo and Valent 2013). Among those, the smut fungus *U. maydis* represents an excellent case to study the infection process. Smut fungi are a large group of biotrophic parasites with currently more than 1500 described species infecting mostly grasses, including important cereal crops such as maize, wheat, barley, and sugar cane (Zuo et al. 2019). The host of *U. maydis* is the sweet corn *Zea mays*, where it can infect all aerial parts of the plant and establishes a biotrophic interface with its host cells.

Biotrophy implies the formation of a tight interaction zone between host and fungal intruder that allows for the exchange of signals and nutrients without initiating apoptosis of host cell tissue. Biotrophic pathogens need to maintain their respective host ‘s viability in order to complete their life cycle. Therefore, *U. maydis* suppresses defense responses, manipulates the metabolism of host cells, and alters their proliferation-rate, ultimately leading to the formation of large spore-filled tumors in the infected tissue (Zuo et al. 2019). The secretion of a variety of effector proteins plays a critical role during this process (Lanver et al. 2017). Effector proteins can be grouped in apoplastic effectors, which remain in the apoplastic space between plant and fungal cells, and cytoplasmic effectors that are further translocated into the host cells’ cytoplasm (Mueller et al. 2008).

This molecular warfare is not restricted to the apoplastic space or the cytosol of host cells. Instead, pathogenic development and tumor formation are accompanied by a thorough remodeling of both plant and fungal cell walls (Matei et al. 2018). These processes support fungal development as the breakdown and import of carbohydrates derived from the host are important sources of carbon for the fungus during growth (Sosso et al. 2019). Sugar sensing and its uptake have thus gained more attention in *U. maydis* in recent years, leading to the identification of several transporters essential for virulence (Wahl et al. 2010; Schuler et al. 2015). The genome of *U. maydis* encodes more than 19 sugar transporters, and most of them are upregulated during pathogenic development (Sosso et al. 2019). Consequently, plants have evolved mechanisms to detect and deplete apoplastic sugar concentrations to hinder fungal growth and activate immune responses (Lemoine et al. 2013; Morkunas and Ratajczak 2014). While these examples are among the first transmembrane proteins studied in the infection context, they also highlight the relevance of membrane-embedded proteins during virulent growth of smut fungi.

However, there is little known on specialized membrane proteins involved in signaling, stimuli recognition, and thus establishing a compatible interaction with the respective host plants. In one case, the membrane protein Pit1, encoded within the pit (**p**rotein **i**mportant for **t**umors) gene cluster, is required for tumor formation (Doehlemann et al. 2011). It has been reported to localize to hyphal tips, although the precise molecular function remains unclear.

Here, we have analyzed a set of six genes showing elevated expression levels during pathogenic development of *U. maydis* (Lanver et al. 2018) encoding proteins that harbor predicted transmembrane helices. Of those two show a strong attenuation in virulence upon deletion of their respective genes. Therefore, we name these proteins Vmp1 and Vmp2 for **v**irulence-associated **m**embrane **p**rotein and present a biochemical characterization giving insights into their molecular architecture and suggesting a potential role during virulence of *U. maydis*.

## 2 Material and Methods

### Molecular cloning of expression plasmids

For the plasmid constructions, standard molecular cloning strategies and techniques were applied (Sambrook J, Fritsch EF 1989). All plasmids and primers used in this study are listed in tables **S1** and **S2**. For the overproduction of the C-terminal domain (CTD) of Vmp2, the plasmid pEMGB1-*vmp2*_CTD_ was generated. The overproduced protein will be fused to the solubility-tag GB1 (56 amino acids), including a hexahistidine tag (Huth et al. 1997). To do so, the region encoding the Vmp2_CTD_ was amplified by PCR from genomic DNA of *U. maydis* SG200 and inserted into the *Nco*I/*Xho*I sites of the vector pEMGB1. For the overproduction of the full-length constructs, the genes encoding Vmp1 and Vmp2 were amplified from genomic DNA of *U. maydis* SG200 without the signal peptide and subsequently ligated into the pEMstX1 vector using *Bsa*I restriction sites. The protein constructs will be fused to a Mistics-tag (110 amino acids), including a hexahistidine tag (Roosild et al. 2005). In both plasmids, a tobacco etch virus (TEV) cleavage site is located between expression tag and cloned gene.

### Strains, growth conditions, and plant infection assays

The *Escherichia coli* strain Dh5α (New England Biolabs) was used for cloning purposes. The *E. coli* strain OverExpress™C43 (DE3) (Sigma-Aldrich) was used to express the full-length constructs of Vmp1 and Vmp2. The *E. coli* strain BL21 (DE3) (Novagen) was used to express the CTD of Vmp2. *E. coli* strains were grown under constant shaking in a temperature-controlled incubator. *Zea mays* cv. Early Golden Bantam (EGB, Urban Farmer, Westfield, IN, USA) was used for infection assays with *Ustilago maydis* and grown in a temperature-controlled greenhouse (light and dark cycles of 14 hours at 28 °C and 10 hours at 20 °C, respectively). *U. maydis* strains used in this study are listed in table **S3**. *U. maydis* strains were grown in YEPS_light_ medium (1 % (w/v) yeast extract, 0.4 % (w/v) peptone and 0.4 % (w/v) sucrose) and subsequently adjusted to an OD_600_ of 1.0 using sterile double-distilled water. For the infection of maize plants 500 µl of *U. maydis* cultures were injected into the stem of 7-day-old maize seedlings using a syringe as described by Kämper and coworkers (Kämper et al. 2006).

### Gene knockout in U. maydis

The plasmid pMS73 was digested with *Acc*65I to integrate the respective sgRNA expression cassette via Gibson Assembly, according to Schuster and coworkers (Schuster et al. 2018). The PCR obtained a double-stranded DNA fragment containing the respective target sequences, scaffold, terminator, and the corresponding overlapping sequences. The fragments were cloned into pMS73 yielding pFA001 and pFA003-pFA007 (**Tab. S1**). The target sequences (**Tab. S2**) were designed using the E-CRISP tool (Heigwer, Kerr, and Boutros 2014). The inserts in all plasmids were validated by sequencing.

### Generation of U. maydis complementation constructs

To generate complementation strains of SG200Δvmp1 and SG200Δvmp2, the constructs pFA511 and pFA512 were generated (**Tab. S1**). Genomic DNA from *U. maydis* SG200 containing promoter and open reading frame (ORF) of the respective gene was amplified by PCR using the primers listed in table **S2**. The amplified fragments were introduced into the *Kpn*I/*Not*I sites of plasmid p123 (Aichinger et al. 2003). Prior to transformation, the plasmids were linearized using the restriction enzyme *Sal*I.

### Generation of U. maydis strains

The genes encoding the six putative transmembrane proteins were disrupted in *U. maydis* SG200 using the CRISPR-Cas9 approach recently described for genetic manipulation of *U. maydis* (Schuster et al. 2016). A donor DNA was supplied during transformation to delete the respective ORF from the genome without further disruption of neighboring genes (**Fig. S2**). Isolated *U. maydis* transformants were confirmed for deleting the respective genes by colony PCR using the primers listed in table **S2** and sequencing (**Fig. S2**). To complement the phenotypes of SG200Δvmp1 and SG200Δvmp2, plasmids pFA511 and pFA512 were integrated into the *ip* locus of SG200. Isolated *U. maydis* transformants were confirmed by Southern-blot analysis to ensure single integration events in the *ip* locus (Keon, White, and Hargreaves 1991).

### Production and purification of soluble Vmp2_CTD_

The CTD of Vmp2 was produced in *E. coli* BL21 (DE3) (Novagen). *E. coli* BL21 (DE3) was transformed with pFA508 to produce Vmp2 _CTD_ fused to an N-terminal GB1 tag including a hexahistidine tag. The protein production was performed in auto-inductive Luria-Miller broth (Roth) containing 1 % (w/v) α-lactose (Roth). The cells were grown for 20 h at 30 °C and 180 rpm. The cultures were harvested by centrifugation (4,000 xg, 15 min, 4°C), resuspended in HEPES buffer (20 mM HEPES, 200 mM NaCl, 20 mM KCl, 40 mM imidazole, pH 8.0), and subsequently disrupted using a microfluidizer (M110-L, Microfluidics). The cell debris was removed by centrifugation (50,000 xg, 20 min, 4 °C). The supernatant was loaded onto Ni-NTA FF-HisTrap columns (GE Healthcare) for affinity purification via the hexahistidine tag. The columns were washed with HEPES buffer (10x column volume) and eluted with HEPES buffer containing 250 mM imidazole. Prior to size exclusion chromatography (SEC), the GB1-tag was cleaved off by adding 0.8 mg purified TEV protease directly to the eluate and incubating under constant rotation at 20 °C for 3 hours. Cleaved His-tagged GB1 and remaining TEV protease were removed via a second Ni-NTA purification after buffer exchange to HEPES buffer containing 40 mM imidazole using an Amicon Ultra-10K centrifugal filter (Merck Millipore). The tag-free protein was subjected to SEC using a Superdex S75 Increase 10/300 column equilibrated in HEPES buffer without imidazole and a pH of 7.5. The peak fractions were analyzed using a standard SDS-PAGE protocol, pooled, and concentrated with Amicon Ultra-10K centrifugal filters.

### Production and Purification of membrane proteins

The plasmids pFA659 and pFA670 encoding full-length Vmp2 and Vmp1 were transformed in *E. coli* OverExpress™C43 (DE3) (Sigma-Aldrich). Transformants were grown in Terrific-Broth medium (24 g/l yeast extract, 20 g/l tryptone, 4 ml/l glycerol, buffered with 10 % phosphate buffer pH 7.4 (0.17 M KH _2_PO_4_, 0.72 M K_2_HPO_4_)) under constant shaking at 180 rpm and 37 °C to an OD_600_ of 0.5 – 0.6. The cultures were then cooled to 20 °C, induced with 0.2 M Isopropyl-β-D-thiogalactopyranosid (IPTG), and incubated for 20 h at 20 °C and 180 rpm. The cultures were harvested by centrifugation (4,000 xg, 15 min, 4 °C), resuspended in Tris-buffer (50 mM Tris-Base, 300 mM NaCl, 40 mM imidazole, pH 8.0), and subsequently disrupted using a microfluidizer (M110-L, Microfluidics). The cell debris was removed by centrifugation (8,000 xg, 20 min, 4 °C) and the supernatant was centrifuged (115,000 xg, 1 h, 4 °C) using a fixed-angle rotor (70 Ti, Beckmann) in an ultracentrifuge (Optima XPN-80, Beckmann). The pellet was resuspended in 10 ml Tris-Buffer using a Dounce-homogenizer (Carl Roth). The homogenized pellet was mixed with 10 ml Tris-Buffer containing either 2 % (w/v) Lauryldimethylamine-N-Oxide (LDAO) or 2 % (w/v) Dodecyl-β-D-maltosid (DDM) for Vmp1 and Vmp2, respectively, and incubated for 2.5 h at 4 °C under constant rotation. The solubilized membrane was again centrifuged (115,000 xg, 1 h, 4 °C). The supernatant was loaded onto 1 ml Ni-NTA FF-HisTrap columns (GE Healthcare) for affinity purification via the hexahistidine tag. The detergent concentration was lowered to 0.1 % (w/v) during the Ni-NTA purification of both proteins. Prior to SEC, the Mistics-tag was cleaved off by adding 0.8 mg purified TEV directly to the eluate and incubating under constant rotation at 20 °C for 3 hours. Cleaved His-tagged Mistics and remaining TEV protease were removed via a second Ni-NTA purification after buffer exchange to Tris buffer containing 40 mM imidazole in an Amicon centrifugal filter (Merck Millipore) with adequate cutoff. The protein was subjected to SEC using a Superdex 200 Increase 10/300 column equilibrated in HEPES-buffer (20 mM HEPES, 200 mM NaCl, 20 mM KCl, pH 7.5) containing either 0.1 % (w/v) LDAO or 0.03 % (w/v) DDM for Vmp1 and Vmp2, respectively. The peak fractions were analyzed using a standard SDS-PAGE protocol, pooled, and concentrated with appropriate Amicon centrifugal filters.

### Multi-angle light scattering (MALS)

Multi-angle light scattering coupled size-exclusion chromatography (SEC-MALS) was performed using an Äkta PURE system (GE Healthcare) with a Superdex 200 Increase 10/300 column attached to a MALS detector 3609 (Postnova Analytics) and a refractive index detector 3150 (Postnova Analytics). The column was equilibrated with 0.2 µm filtered HEPES buffer (20 mM HEPES, 200 mM NaCl, 20 mM KCl, pH 7.5) containing either 0.1 % (w/v) LDAO or 0.03 % (w/v) DDM for Vmp1 and Vmp2, respectively. For each measurement, 100 µl of a 50 µM protein solution was injected.

### Mass photometry (MP)

Mass photometry experiments were performed using a OneMP mass photometer (Refeyn Ltd, Oxford, UK). Data acquisition was performed using AcquireMP (Refeyn Ltd. v2.3). Mass photometry movies were recorded at 1 kHz, with exposure times varying between 0.6 and 0.9 ms, adjusted to maximize camera counts while avoiding saturation. Microscope slides (70 x 26 mm) were cleaned 5 minutes in 50 % (v/v) isopropanol (HPLC grade in Milli-Q H_2_O) and pure Milli-Q H_2_O, followed by drying with a pressurized air stream. Silicon gaskets to hold the sample drops were cleaned in the same manner fixed to clean glass slides immediately prior to measurement. The instrument was calibrated using NativeMark Protein Standard (Thermo Fisher) immediately prior to measurements. Immediately prior to mass photometry measurements, protein stocks were diluted directly in HEPES buffer. Typical working concentrations of Vmp1 and Vmp2 were 25-50 nM for the actual measurement. Each protein was measured in a new gasket well (i.e., each well was used once). To find focus, 18 µl of fresh room temperature buffer was pipetted into a well, the focal position was identified and locked using the autofocus function of the instrument. For each acquisition, 2 µL of diluted protein was added to the well and thoroughly mixed. The data were analyzed using the DiscoverMP software.

### Confocal light microscopy

The proliferation of *U. maydis* in infected maize leaf tissue was visualized by confocal microscopy as described previously (Tanaka et al. 2014). A leaf area of 1 cm ^2^ located 2 cm below the injection site was excised 2 days post-infection (dpi). The leaf samples were destained with ethanol and treated with 10 % (w/v) potassium hydroxide at 85°C for 4 h. The fungal hyphae were stained with Wheat Germ Agglutinin-Alexa Fluor 488 (WGA-AF488, Invitrogen). The plant cell walls were stained with propidium iodide (Sigma-Aldrich) by incubating decolorized samples in staining solution (1 µg/ml propidium iodide, 10 µg ml−1 WGA-AF488) and observed with a TCS-SP8 confocal laser-scanning microscope (Leica Microsystems) under the following conditions: WGA-AF488: excitation at 488 nm and detection at 500 –540 nm; propidium iodide: excitation at 561 nm and detection at 580–660 nm.

### Fungal stress assays

Fungal strains were grown in YEPS_light_ medium (1 % (w/v) yeast extract, 0.4 % (w/v) peptone and 0.4 % (w/v) sucrose) to an OD_600_ of 1.0. The cells were pelleted and resuspended in sterile double distilled H_2_O to an OD_600_0.1. For the induction of filament formation, 10 µ l of serial dilutions were spotted on potato-dextrose charcoal plates (Holliday 1974). The stress assays were performed on CM plates (Holliday 1974) supplemented with 750 µM calcufluor white (Sigma-Aldrich), 3 mM hydrogen peroxide (H_2_O_2_), 1 M NaCl or 1 M sorbitol. Images were taken after overnight incubation at 28 °C.

### Statistical analysis

Disease symptoms of infected plants were scored at 12 dpi using the previously established scoring scheme by Kämper and colleagues (Kämper et al. 2006). Disease symptoms were quantified based on three biological replicates and are presented as stacked histograms. Significant differences among disease symptoms within individual disease categories were determined by Student’s t-test. The raw data of all infection assays and the statistical analysis can be found table **S5**.

### Accession numbers

The genes and encoding protein sequences from *U. maydis* are available at NCBI under the following accession numbers.: *vmp1* (UMAG_00032), XP_011386009.1; *vmp2* (UMAG_01689), XP_011387666.1; UMAG_01713, XP_011387687.1; UMAG_04185, XP_011390672.1; UMAG_10491, XP_011390314.1; UMAG*_*03474, XP_011389930.1.

## 3 Results

### Identification of membrane proteins critical for pathogenic development of U. maydis

To identify membrane proteins that show an increase in transcript abundance during infection stages associated with biotrophic development of *Ustilago maydis*, we analyzed the transcriptomic data obtained by Lanver and coworkers (Lanver et al. 2018). Highly upregulated protein-encoding genes were then examined for the presence of potential transmembrane helices (TMs) using the Consensus Constrained TOPology prediction web server CCTOP (**Fig. S1**) (Dobson, Reményi, and Tusnády 2015). By this approach, we could identify six genes strongly elevated during infection and their respective proteins containing at least one predicted TM. They show their strongest expression two to four days post-infection while not induced in axenic culture under non-infective conditions (**Fig. 1A**). These proteins are UMAG_00032, UMAG_01689, UMAG_01713, UMAG_0 3474, UMAG_04185, and UMAG_10491.

**Figure 1.**
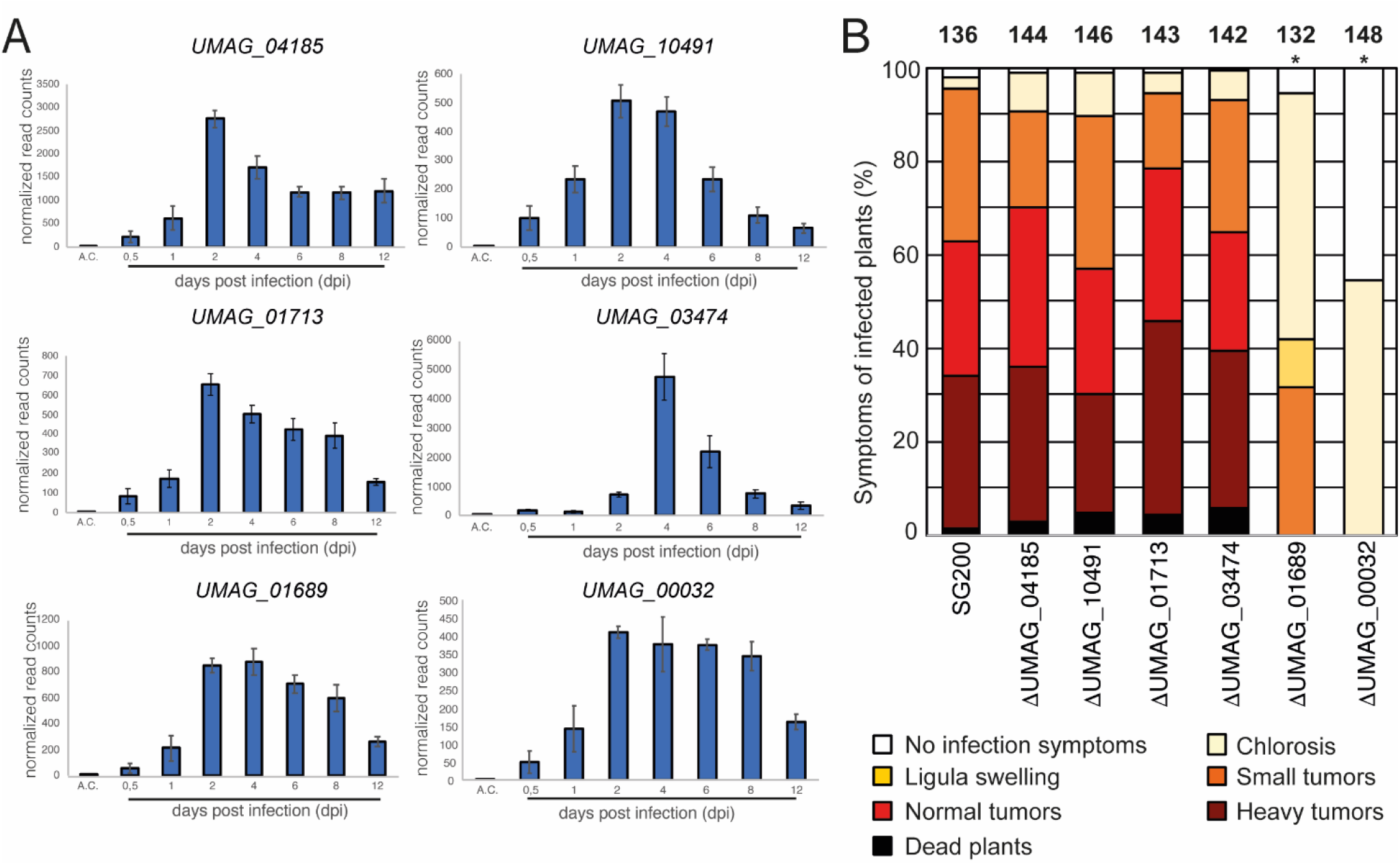
Identification of a transmembrane protein important for virulence. **A**. The expression pattern of genes encoding transmembrane proteins in *U. maydis* during plant infection re-analyzed from RNA sequencing data (Lanver et al. 2018). A.C., expression level in axenic culture. The numbers below the bars indicate the days post inoculation (dpi). Error bars indicate ± SD. **B**. Virulence assay of genes encoding transmembrane proteins in the *U. maydis* SG200 background. Disease symptoms were quantified on maize leaves 12 days post infection. Similar results were observed in three independent experiments. Shown is the mean percentage of plants placed in a particular disease category. The number of infect ed plants is indicated above the bars. The asterisk indicates a significant difference in infection symptoms between SG200, SG200Δvmp1 and SG200Δvmp2.

To evaluate these protein’s impact on virulence, we deleted their respective genes in the solopathogenic *U. maydis* strain SG200 (Kämper et al. 2006). The gene deletion was performed using a CRISPR-Cas9-based approach as described by Schuster and coworkers (Schuster et al. 2016). A donor DNA was supplied to delete the respective open reading frames (ORF’s) from the genome while keeping the surrounding genetic environment intact (**Fig. S2**). The deletion of four genes resulted in a wildtype-like behavior during maize infection experiments (*UMAG_01713, UMAG_03474, UMAG_04185*, and *UMAG_10491*), the two other genes (*UMAG_00032* and *UMAG_01689*) lead to attenuation in virulence (**Fig. 1B**). To investigate whether the differences in phenotypical symptoms between the deletion strains of *UMAG_00032, UMAG_01689*, and SG200 are significant, we scored the disease symptoms of each infected plant using a previously established scoring scheme (Kämper et al. 2006). For the significance-analysis, we performed a two-sided Student’s t-test. Our analysis for each category confirmed that the differences are significant, with p-values below 0.05 for several categories (**Tab. S5**).

Our results reveal two TM proteins that strongly impact the virulence of *U. maydis* during maize infection. Therefore, we named both genes *vmp1* (*UMAG_00032*) and *vmp2* (*UMAG_01689*) for **v**irulence-associated **m**embrane **p**rotein 1 and 2.

### Vmp1 and vmp2 are conserved among related smut species

*Vmp1* encodes a protein of 142 amino acids (aa), whereas *vmp2* encodes a 335 aa long protein. Both proteins contain an N-terminal signal peptide (SP) of 25 aa, as predicted by SignalP-5.0 (Almagro Armenteros et al. 2019). Our *in silico* analyses indicate that both proteins harbor one TM helix spanning the residues 60 to 77 in Vmp1 and residues 100 to 115 in Vmp2 (**Fig. S1)**. The N-terminal domain (NTD) of both proteins is predicted to be extracellular (**Fig S1**).

In a next step, we analyzed the genetic context of both proteins in *U. maydis* and compared it to related smut fungi. Using the Basic Local Alignment Search Tool (BLAST) we identified Vmp1 orthologs in the genomes of *Pseudozyma hubeiensis* SY62, *Kalmanozyma brasiliensis* GHG001, *Sporisorium reilianum* SRZ2, *Ustilago trichophora, Sporisorium scitamineum, Moesziomyces antarcticus, Moesziomyces aphidis* DSM 70725, and *Testicularia cyperi* with identities ranging from 58 % to 34 % (determined by CLUSTAL2.1) (**Fig. S3A**). However, it was absent in *Ustilago hordei* or *Ustilago bromivora* with the genetic context being similar to *U. maydis* (**Fig. 2A**). A protein related to Vmp1 was also identified in the genome of *T. cyperi* a pathogen of *Rhynchospora* spp. (Kijpornyongpan et al. 2018). The genetic context showed differences to the closely related species due to the ancestral nature of *T. cyperi* (**Fig. 2A**).

**Figure 2.**
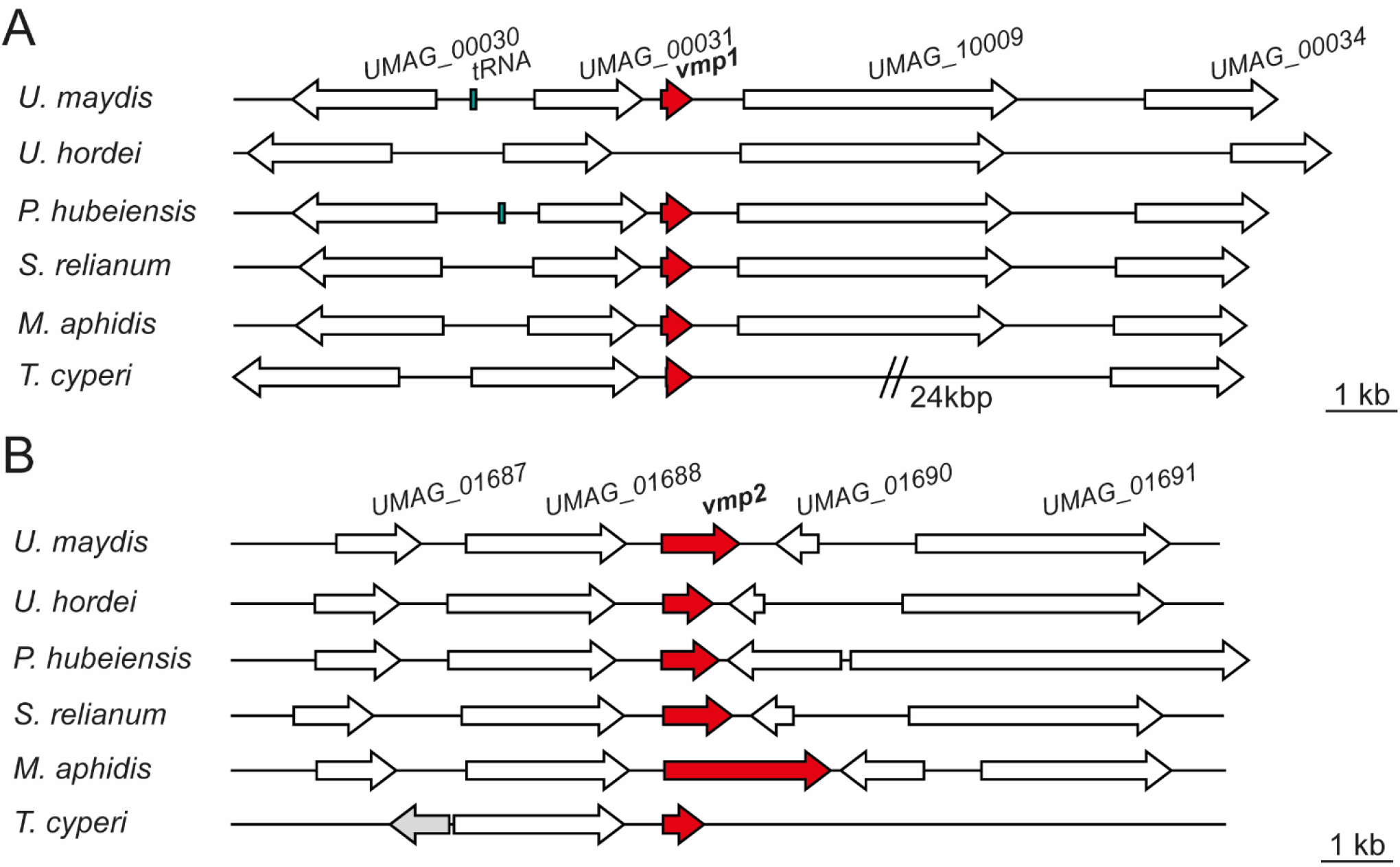
Vmp1 and Vmp2 orthologs are conserved in related smut fungi. Schematic picture of gene loci encoding Vmp1 (**A**) and Vmp2 (**B**) and orthologs in the related smut pathogens *Ustilago hordei, Pseudozyma hubeiensis, Sporisorium relianum, Moesziomyces aphidis* and *Testicularia cyperi*. White arrows indicate genes found in all of the respective species, while t he grey gene was solely present in the genome of *T. cyperi*.

The neighboring genes encode a proline dehydrogenase (*UMAG_00030*), a TM protein of unknown function (*UMAG_00031*), a Zn_2_-C6 fungal-type transcription factor (*UMAG_10009*), and a putative MFS transporter (*UMAG_00034*) (**Fig. 2A**). These genes are also induced during axenic growth and might thus not be directly related to virulence. However, *UMAG_00034* shows elevated transcript levels between 24 h and 48 h post-infection while not induced during axenic growth (Lanver et al. 2018).

We also identified orthologs of Vmp2 in a variety of related smut fungi (**Fig. S3B**). Namely, *P. hubeiensis* SY62, *U. bromivora, Sporisorium graminicola, S. reilianum* SRZ2, *U. hordei, K. brasiliensis* GHG001, *U. trichophora, M. antarcticus, S. scitamineum*, and *T. cyperi*. Here, the sequence identities ranged from 43 % to 36 % (**Fig. S3B**). Notably, Vmp2 is highly conserved from amino acid 82 to 195 (within the Vmp2 sequence from *U. maydis*), while the C-terminus shows a higher degree of deviation in the investigated orthologs (**Fig. S3B**). In Ustilaginaceae, the loci of *vmp2* are similarly to *vmp1* highly syntenic although the intergenic region towards *UMAG_01690* and its orthologs shows some length differences (**Fig. 2B**). The neigbhouring genes include an OBG-type G-domain-containing protein (*UMAG_01687*), a putative nuclear transport factor (*UMAG_01688*), a secreted effector protein of unknown function (*UMAG_01690*) and a DNA helicase (*UMAG_01691*) (**Fig. 2B**).

### Vmp1 hinders fungal infection after penetration of the plant epidermis

The *vmp1* deletion strain showed the strongest reduction in virulence with tumor formation being entirely abolished in infected plants (**Fig. 3A, B**). Anthocyanin production was observed in the vicinity of the infection site, a universal sign of infections and thus the presence of infectious hyphae (Tanaka et al. 2014). The deletion strain SG200Δvmp1 was complemented by integrating a single copy of *vmp1* into the *ip* locus (SG200Δvmp1-vmp1, **Fig. 3A**). The complementation did not fully restore SG200Δvmp1, leading mainly to the formation of smaller tumors and larger ones only to a lesser extent (**Fig. 3A, B**). Thus, we wanted to know whether SG200Δvmp1 remains able to grow inside vascular bundles and elicits a plant defense response or whether fungal growth is arrested after penetration of the epidermal layer.

**Figure 3.**
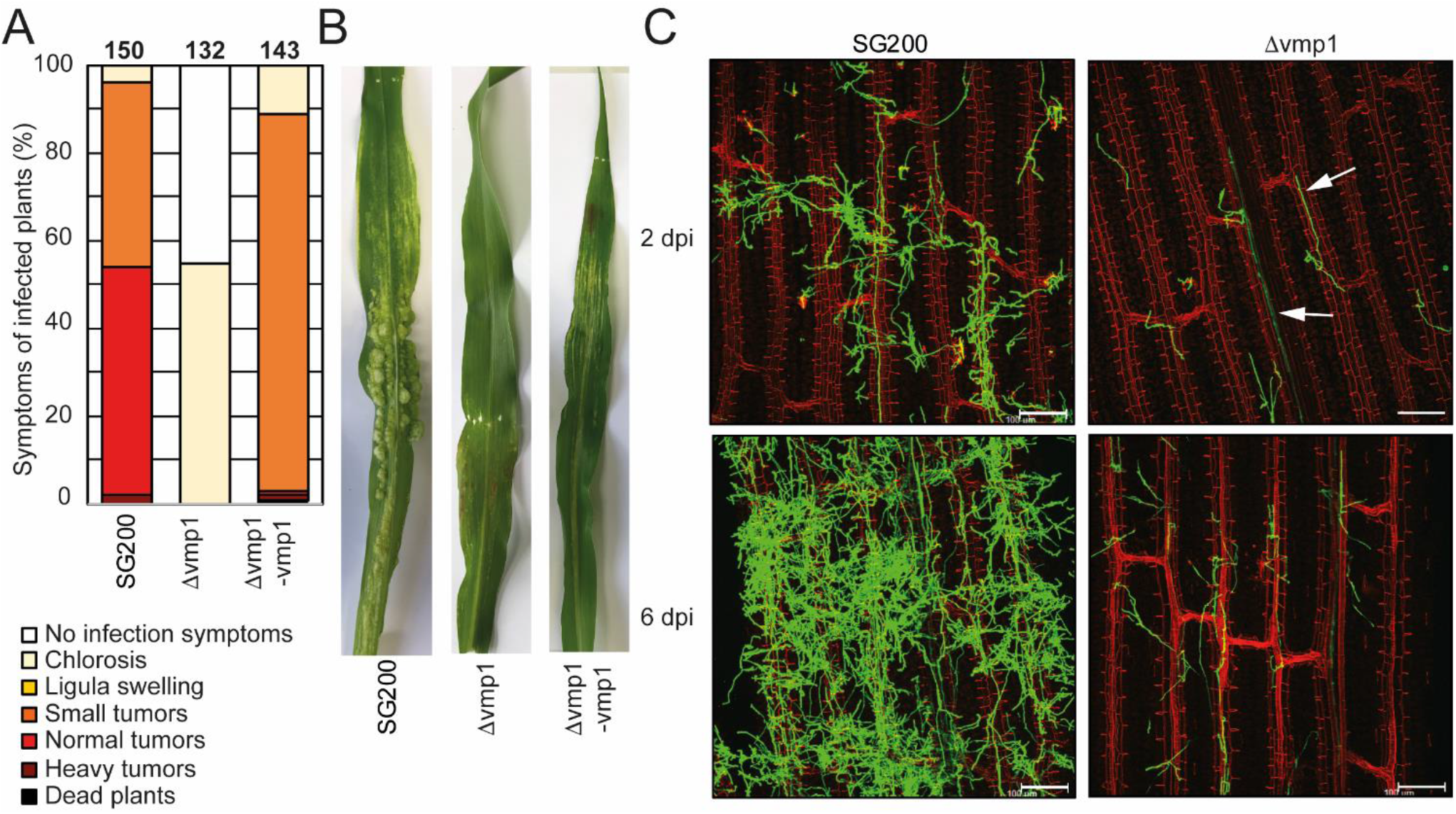
Vmp1 is required for virulence. **A**. Virulence assay of the SG200Δvmp1 mutant strain and SG200Δvmp1-vmp1 complementation strain in an *U. maydis* SG200 background. The mean percentage of disease symptoms in the different categories is shown that were quantified based on three biological replicates. The number of infected plants is indicated above the bars. **B**. Macroscopic pictures of maize leaves 12 days post infection with *U. maydis* SG200, SG200Δvmp1 and SG200Δvmp1-vmp1. **C**. Leaf tissues infected with SG200 and SG200Δvmp1 were stained with WGA-AF488 and propidium iodide at 2 and 6 dpi. Green color indicates fungal hyphae and red color indicates leaf vascular bundles. Bar = 100 μm.

To detect differences in host colonization, we visualized fungal hyphae by staining with WGA-AF488 at 2 and 6 days post-infection (dpi) (**Fig. 3C**). It became apparent that SG200Δvmp1 has a reduced number of fungal hyphae on the plant leaf surface combined with less proliferation (**Fig. 3C**). However, hyphae could still penetrate the epidermal layer and grow inside the vascular bundles (**Fig. 3C**). Fungal growth was seemingly arrested at this stage as the amount of fungal material inside the plant leaves was not drastically increased at 6 dpi (**Fig. 3C**). To rule out that the reduced virulence was due to reduced growth and stress sensitivity, we grew SG200Δvmp1 in the presence of NaCl, sorbitol, calcofluor white, and H_2_O_2_. However, mutant strains were indistinguishable from SG200 (**Fig. S4**).

In conclusion, we show that the TM protein encoded by *vmp1* is essential for full virulence and might be important for establishing the biotrophic interface. It is conserved among related smut fungi (**Fig. S3A**) indicating that its function might also be conserved among these relatives.

### Vmp2 leads to reduced tumor formation

The deletion of *vmp2* led to a strong reduction in virulence of *U. maydis*, with solely small tumors being formed (**Fig. 4A**). We complemented SG200Δvmp2 by integrating a single copy of *vmp2* into the *ip* locus (SG200Δvmp2-vmp2, **Fig. 4A, B**). This complementation could fully restore the phenotype of SG200Δvmp2. To rule out that the deletion of *vmp2* leads to altered growth of *U. maydis* under stress conditions, we grew SG200Δvmp2 in the presence of NaCl, sorbitol, calcofluor white and H_2_O_2_ and did not detect differences from SG200 (**Fig. S4**).

**Figure 4.**
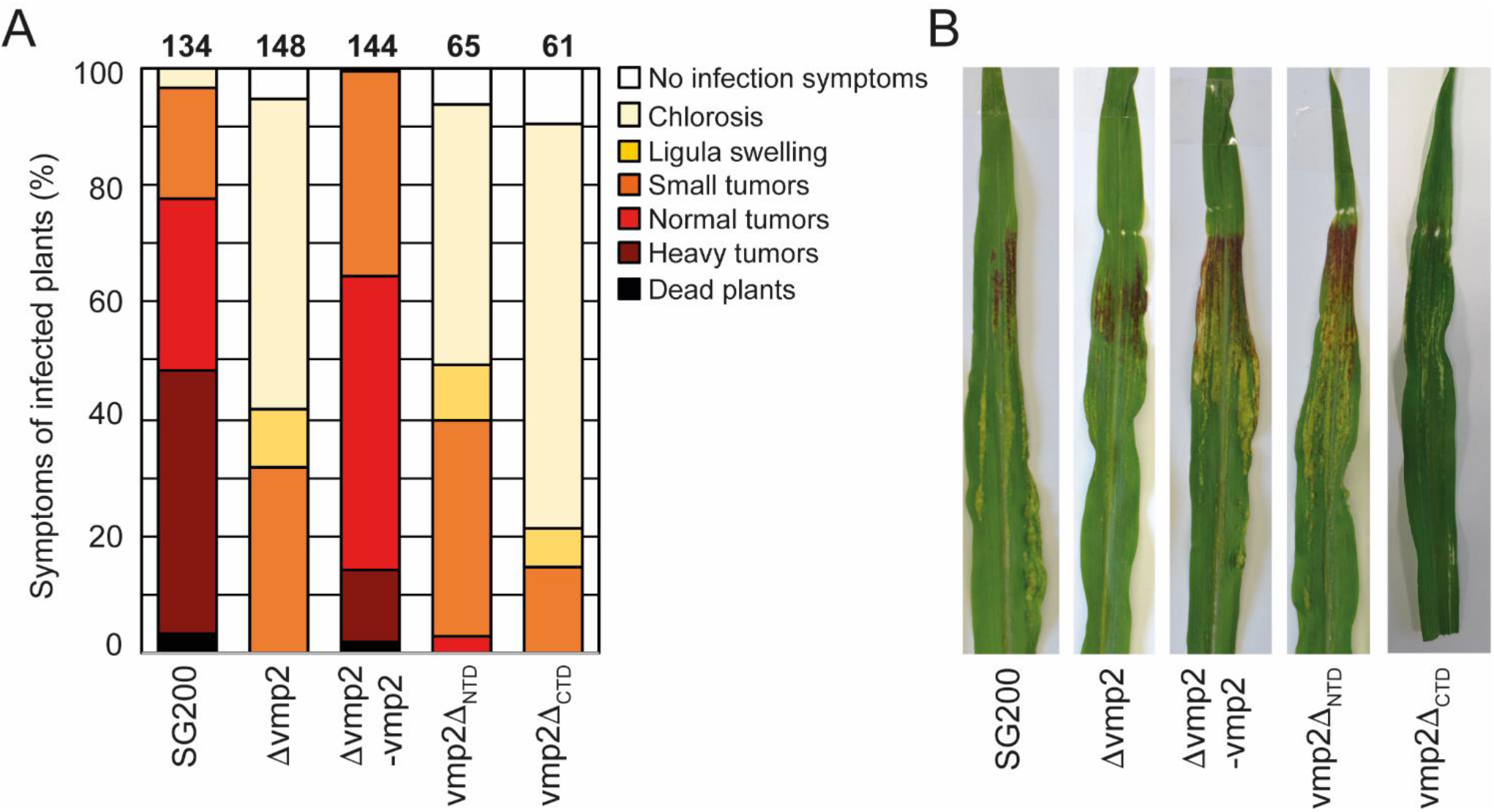
Vmp2 is required for virulence. **A**. Virulence assay of the SG200Δvmp2 mutant strain and SG200Δvmp2-vmp2 complementation strain in *U. maydis* SG200 background. Disease symptoms were quantified based on three biologica l replicates. The number of infected plants is indicated above the bars. **B**. Macroscopic pictures of maize leaves infected by U. maydis SG200, SG200Δvmp2 and SG200Δvmp2-vmp2 at 12 dpi.

In the next step, we aimed to understand how deleting the two predicted soluble domains would impact the function of Vmp2 *in vivo* (**Fig. S1**). We generated two constructs deleting either the predicted extracellular NTD or the cytosolic CTD and transformed *U. maydis* SG200 to perform infection assays (**Fig. S2D**). Our experiments show that SG200vmp2_ΔCTD_phenocopies SG200Δvmp2, while SG200vmp2_ΔNTD_ is less attenuated in virulence (**Fig. 4A, B**).

Taken together, we can show that *vmp2* is an essential player for the infection process in *U. maydis* and potentially related organisms. Additionally, our infection experiments indicate that the CTD of Vmp2 is important for full virulence.

### Vmp1 shows concentration-dependent oligomerization

To allow for a biochemical investigation of Vmp1, we cloned the open reading frame without the signal peptide (residues 1-20) for heterologous protein production in *E. coli* (see materials & methods). First expression and solubility tests did not allow to purify the full-length protein in amounts sufficient for biochemical analysis. Thus, we generated a construct that includes an N-terminal Mistics-tag (MstX) separated by a TEV protease cleavage site. This 110 amino acid long protein tag forms four transmembrane helices and inserts autonomously in the membrane. It has been used to improve the expression of membrane proteins in several cases (Roosild et al. 2005). In our case, the production of MstX-Vmp1 was drastically enhanced compared to protein production without the fusion-tag. Attempts to solubilize MstX-Vmp1 from the membrane fraction using Dodecyl-β-D-maltosid (DDM) failed and thus we tested a variety of commercially available detergents. Solubilization was only achieved employing Lauryldimethylamine-N-Oxide (LDAO). Notably, all attempts to cleave the MstX tag via TEV cleavage only resulted in inefficient and partial cleavage. It is likely that the spacing between the membrane-embedded MstX and the membrane spanning helix within Vmp1 (residues 60 – 77) might not allow for a proper TEV recognition and cleavage. Consequently, we used the full-length fusion protein for biochemical analysis.

Purified MstX-Vmp1 was subjected to size-exclusion chromatography coupled multi-angle light scattering (SEC-MALS) using a Superdex 200 Increase 3.2/300 column equilibrated with SEC buffer including 0.1% LDAO (see materials & methods). The protein eluted in a single peak at 1.62 ml corresponding to 90 kDa according to the calibration calculation for this column (**Fig. 5A**). Our analysis with MALS and refractive index resulted in a mass of 113 ± 17 kDa and thus yielded a slightly higher molecular weight (**Fig. 5A**). The calculated mass of the MstX-Vmp1 fusion protein is around 31 kDa. MALS allowed us to clearly distinguish between empty micelles and the membrane protein-detergent complexes. Notably, the molecular weight of free LDAO micelles was found to be 40 ± 5 kDa in our experiments and thus a bit larger than 16-20 kDa reported in literature (Timmins et al. 1988). As membrane proteins are likely not embedded into detergent micelles but rather form membrane protein-detergent complexes (Chaptal et al. 2017), our results indicate that two or three Vmp1 molecules would be encaged by LDAO detergent molecules.

**Figure 5.**
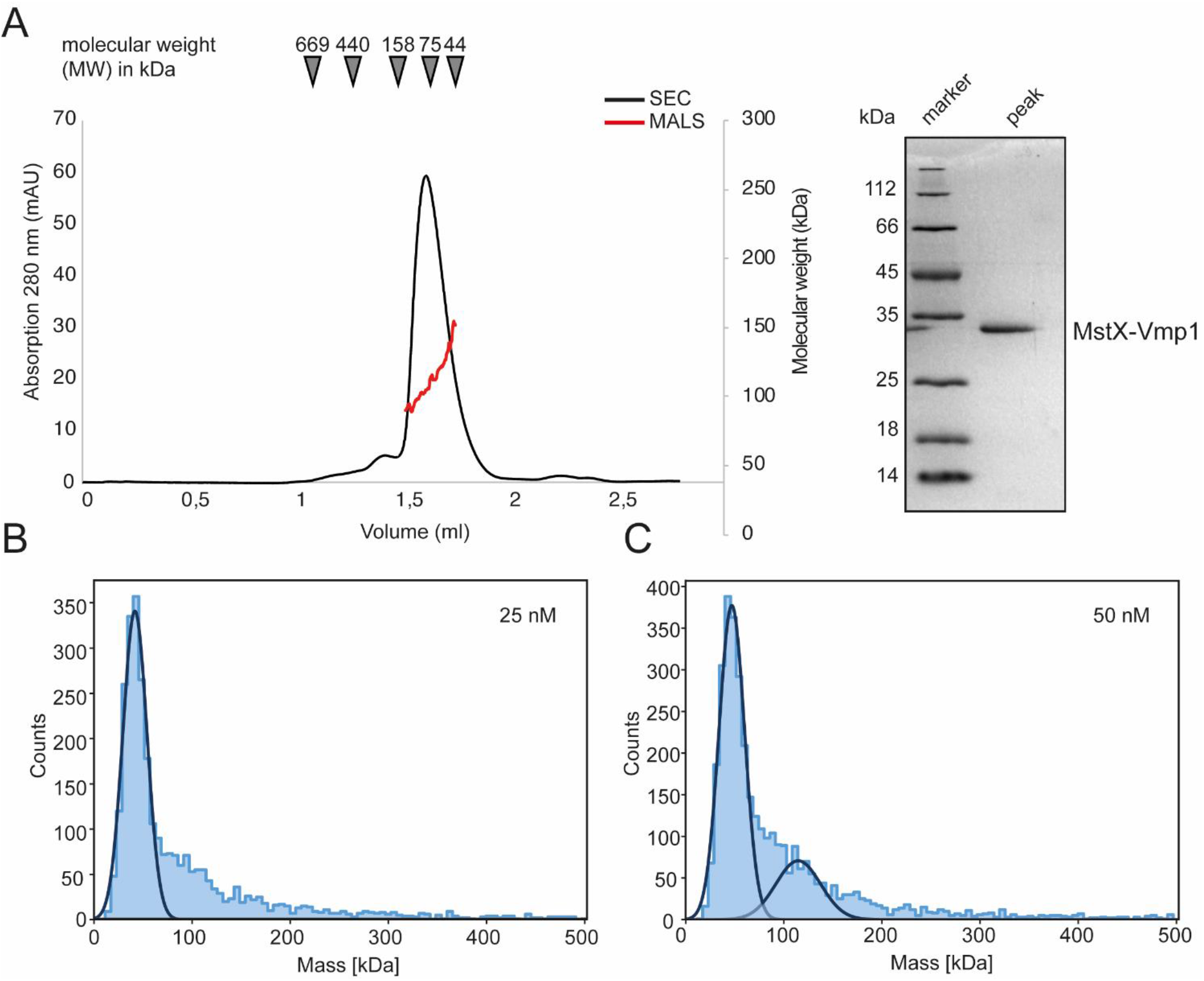
Biochemical analysis of Vmp1. **A**. Multi-angle-light scattering coupled size-exclusion chromatography (SEC-MALS) of full length MstX-Vmp1. The black line depicts the absorption at 280 nm, while the red line corresponds to the molecular weight as determined by MALS. The inset shows a SDS PAGE of the peak fraction. **B** and **C**. Mass photometry of Vmp2 in 0.1% LDAO at 25 and 50 nM concentration, respectively.

To achieve a better resolution of Vmp1 oligomerization, we employed mass photometry, a method that became recently available and allows rapid and reliable determination of the dynamic molecular weight of macromolecules in solution (Olerinyova et al. 2021; F et al. 2020). We firstly used a final concentration of 25 nM MstX-Vmp1 for mass photometric analysis which was achieved by rapid 1:10 dilution of a 250 nM solution into SEC buffer without detergent. Approximately 60 % of MstX-Vmp1 had a measured mass of 42 kDa (**Fig. 5B**) suggesting a monomer of MstX-Vmp1 and ∼50 LDAO detergent molecules (11 kDA). A subfraction higher molecular weight assemblies was also visible, however gaussian fitting was not possible at this concentration. When using 50 nM of MstX-Vmp1, a second gaussain could be fitted additionally to the 60 % of molecules with a mass of 42 kDa indicating the presence of a 118 kDa species containing 20 % of all molecules (**Fig. 5C**). To rule out that no empty LDAO micelles were detected, we subjected a buffer containing no protein and only LDAO at the working concentration of 0.01% to mass photometry. However, no events were detectable suggesting that micelles are not formed at this detergent concentration.

Taken together, we conclude that Vmp1 mainly occurs as a monomer but forms higher oligomeric species at higher concentrations.

### Vmp2 is a dimeric membrane protein

In a next step, we aimed to investigate Vmp2 after heterologous protein production in *E. coli*. Similar to Vmp1, the expression of full-length Vmp2 was insufficient for biochemical analyses and only the fusion of an N-terminal Mistics-tag allowed to obtain adequate amounts of membrane-bound protein. Vmp2 could be solubilized with DDM and was purified using a Superose 6 Increase 10/300 column (GE Healthcare) equilibrated with SEC buffer and 0.03 % DDM (see materials & methods). The protein eluted at 17.22 ml corresponding to a molecular weight of approx. 83 kDa (**Fig. 6A**).

**Figure 6.**
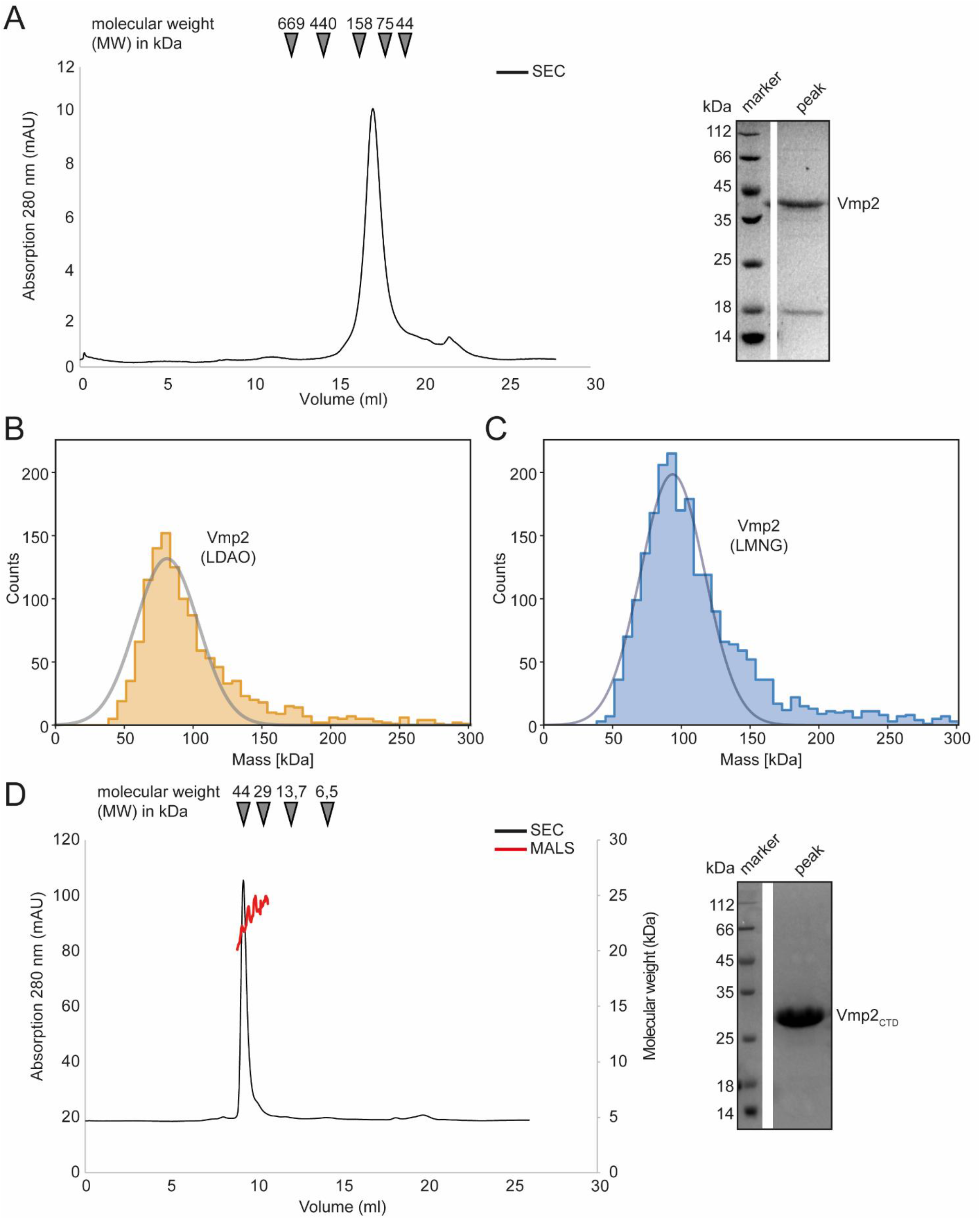
Biochemical analysis of Vmp2. **A**. SEC chromatogram of full length Vmp2. The inset shows a SDS PAGE of the peak fraction. **B**. Mass photometry of Vmp2 in 0.1% LDAO. **C**. Mass photometry of Vmp2 in 0.001% LMNG. **D**. Multi-angle-light scattering coupled size-exclusion chromatography (SEC-MALS) shows that the C-terminal domain (CTD) of Vmp2 (aa 120 – 335) is monomeric with an apparent molecular weight (MW) of 24 kDa. The black line depicts the absorption at 280 nm, while the red line corresponds to the molecular weight as determined by MALS. The inset shows a SDS PAGE of the peak fraction.

We again employed mass photometry to accurately determine the molecular weight of Vmp2 and investigate whether different oligomeric species might be visible even at nanomolar concentrations. However, using DDM as detergent, 0.03 % (w/v), which is aquivalent to 600 µM and thus generated a strong detergent background that did not allow us to distinguish between empty micelles and Vmp2. Thus, we investigated whether Vmp2 would be stable in LDAO or lauryl maltose neopentyl glycol (LMNG), a detergent that contains two DDM moieties and has a very low CMC at 10 µM which is perfectly suited for mass photometry. We thus solubilized Vmp2 using DDM and exchanged the detergent during Ni-ion affinity chromatography and applied the protein to a Superose 6 Increase 10/300 column equilibrated in 0.1 % LDAO or 0.001 % LMNG, respectively.

Firstly, Vmp2 purified in the presence of 0.1 % LDAO was measured (**Fig. 6B**). To remove excess detergent micelles during mass photometry, a stock solution at 1 µM of Vmp2 was rapidly diluted 1:10 in SEC buffer without detergent. A Gaussian fit of the peak fraction contained 92 % of all measured molecules at a MW of 81 kDa. In a second approach, we used Vmp2 solubilized in 0.001 % LMNG and again rapidly diluted it 1:10 in SEC buffer containing no detergent. Here, we could fit 84 % of all counts resulting in a MW of approximately 94 kDa (**Fig. 6C**). The mass differences between the LDAO and LMNG solubilized Vmp2 likely is a result from the different protein-detergent complex sizes formed by the two detergent molecules. As Vmp2 has a theoretical molecular weight of 32 kDa, the 81 kDa would correspond to a dimer of Vmp2 and ∼75 LDAO (17 kDa) detergent molecules, while the 94 kDa suggest a Vmp2 dimer and ∼30 LMNG (30 kDa) detergent molecules.

In summary, our mass photometry results are in agreement with the MW calculated from size exclusion chromatography and indicate the presence of a Vmp2 dimer.

### The CTD of Vmp2 is largely unstructured and does not contribute to dimerization

Next, we investigated the predicted cytosolic CTD of Vmp2. We subjected purified Vmp2 _CTD_ to a Superdex 75 Increase 10/300 column. The protein eluted at 9.28 ml which corresponds to a molecular weight of 45 kDa (**Fig. 6D**). However, multi-angle light scattering coupled SEC (SEC-MALS) unambiguously revealed a MW of 25 ± 1,5 kDa of Vmp2 _CTD_ (**Fig. 5D**). Our secondary structure and disorder prediction through PSIPRED indicated that residues 200 to 335 are potentially disordered (**Fig. S5**). As disordered or non-globular proteins show a different migration behavior than the SEC-standard, this would explain the discrepancy between SEC and MALS MW calculation. In conclusion, we can show that Vmp2 is dimeric membrane protein with a CTD that is largely unstructured and does not contribute to dimerization.

## 4 Discussion

In this study, we have identified six genes that are strongly induced between 0.5 and 2 days post-infection (dpi) and remain upregulated until 12 dpi (**Fig. 1A**), while not being expressed in axenic culture. This expression pattern correlates with establishing and maintaining biotrophy, a critical feature of pathogenic development in smut fungi (Lanver et al. 2018). Our *in silico* analysis suggested that all of them harbor at least one transmembrane spanning helix, rendering them interesting targets as proteins associated with virulence in smut fungi are predominantly soluble effectors (Lanver et al. 2017). The deletion of two of them, subsequently named Vmp1 and Vmp2 (**v**irulence associated **m**embrane **p**rotein), resulted in a strong attenuation of virulence during maize infection, while growth of the deletion strains was neither affected in axenic liquid culture nor in the presence of various stress causing agents (**Fig. S4**). We can thus conclude that both Vmp1 and Vmp2 are important during pathogenic but not axenic growth of *U. maydis*. Attempts to reveal a potential function of these TM proteins by the prediction of functional domains yielded no results for Vmp1 and Vmp2 using the DomPred server embedded in the PSIPRED algorithm (Buchan and Jones 2019).

To shed light on the function of Vmp1, we inspected the deletion strains in more detail. Deletion of Vmp1 led to a strong attenuation of fungal growth that was arrested after epidermal penetration (**Fig. 2C**) although some hyphae were still visible growing inside vascular bundles. Notably, tumor formation on maize leaves inoculated with vmp1 mutant strains was not observed in infection experiments. Vmp1 thus plays a critical role during the early infection stages. Notably, vmp1 mutant strains still elicited a plant defense response as anthocyanin production could still be observed on infected plant leaves.

Our biochemical analysis suggested that Vmp1 predominantly occurs as a monomer (**Fig. 5B**) as the cellular concentrations of Vmp1 will most likely be low. This is further supported by the gene expression data as *vmp1* shows the lowest expression of all six transmembrane protein encoding genes investigated (**Fig. 1A**). During investigation of the genomic context of *vmp1*, it became apparent that the gene *UMAG_00031* is found in the same orientation upstream of vmp1 in several related species. A recent study demonstrated that *UMAG_00031* encodes a putative transmembrane protein potentially involved in pH regulation (Cervantes-Montelongo et al. 2020). In contrast to SG200Δ*vmp1*, UMAG_00031 mutant strains showed reduced growth under pH stresses as well as in the presence of sorbitol and NaCl (Cervantes-Montelongo et al. 2020). The study suggested UMAG_00031 to be a member of the Pal/Rim pathway in *U. maydis*, a widely conserved signaling pathway involved in pH adaptation (Selvig and Alspaugh 2011; Fonseca-García, León-Ramírez, and Ruiz-Herrera 2012). However, our data indicate that Vmp1 is most likely not directly involved in pH adaptation or regulation. It might still play an accessory role in these processes serving e.g. as adaptor protein. Here, future research might identify a connection towards pH related regulation to during plant infection.

Vmp2 (UMAG_01689) has already been identified to contribute to virulence in *U. maydis* (Uhse et al. 2018). In their study, the authors also showed that the fungal biomass is strongly reduced in infected plant leaves. However, as the knockout was only delivered as a proof-of-concept of their method to identify genes essential for virulence, no further information on Vmp2 was provided. Our data confirm the phenotype observed by Uhse and coworkers (**Fig. 4A**). Furthermore, we can show that Vmp2 has a short N-terminal (NTD) and a long C-terminal domain (CTD). While deletion of the CTD phenocopies SG200Dvmp2, strains deleted for the NTD cause slightly more severe symptoms on infected plants. This suggests that the CTD is indispensable for virulence. Sequence alignments to homologs from other smut fungi show that the C-termini is highly variable, while the region surrounding the membrane spanning helix is conserved (**Fig. S3**). Our analysis by SEC-coupled MALS confirmed that the CTD is largely unstructured. Proteins containing unstructured regions have been characterized in the context of many scenarios and can make up substantial amounts of the total protein content (Van Der Lee et al. 2014). A possible scenario is that the unstructured region of Vmp1 becomes ordered in the context of an interaction partner. Here, the sequence variability in related organisms suggests that this interface is species-specific. Another possible explanation might be that the unstructured domain is involved in membrane shaping or impacts the local membrane heterogeneity (Fakhree, Blum, and Claessens 2019). A thorough investigation of the interactome of Vmp2 *in planta* might deliver an explanation for the role of CTD of Vmp2 during maize infection of *U. maydis*.

In conclusion, we here present two membrane proteins that act as virulence factors during maize colonization of *U. maydis*. While we deliver an initial characterization of the two proteins expanding the current knowledge on virulence associated membrane proteins of smut fungi, future research needs to address their precise functions.

## Supporting information

Weiland and Altegoer Supplement Final

## 6 Author Contributions

F.A. conceived of the project and designed the study. F.A. and P.W. performed experiments, analysed data and wrote the paper.

## 8 Acknowledgments

We thank Gert Bange for financial support and helpful discussion on the manuscript and Regine Kahmann for her continuous support throughout the work on this manuscript. We are grateful to Georg Hochberg for access to the Refeyn One mass photometer and to Pietro Giammarinaro for subsequent assistance with data collection.

## 9 Contribution to the Field Statement

Fungal phytopathogens are an increasing threat to important crops such as wheat, sugar cane, and maize. The smut fungi *Ustilago maydis* specifically infects the sweet corn *Zea mays* and leads to annual crop losses of up to 20 %. The pathogen needs to establish a biotrophic interaction with its host in order to gain access to valuable resources for life cycle completion. Maintaining this tight interaction, without triggering any immune responses of the host is mainly accomplished by the secretion of a variety of effector proteins in the apoplastic space between host and parasite. Many of those soluble effector proteins have recently been described to be involved in the attenuation of defense mechanisms and the manipulation of host cell metabolism. However, we still don’t fully understand the underlying principles of important processes such as cellular host recognition and attachment, the endosomal transport of proteins to the apoplastic space and host cells, or the uptake of signals and nutrients. Membrane-bound proteins with elevated expression during the pathogenic development of *U. maydis* are highly likely to be involved in these processes and should therefore attract more attention in this research field. By delivering an initial characterization of two membrane proteins required for virulence, we highlight the importance of membrane proteins for understanding the infection process of *U. maydis*.

