## Supplementary material for "Identification and characterization of two transmembrane proteins required for virulence of *Ustilago maydis*": Weiland and Altegoer Supplement Final

**Supplementary information**

**Table S1.** Plasmids used in this study.

| **Plasmid** | **Usage** | **Citation** |
| --- | --- | --- |
| pEMGB1 | Protein overexpression with N-terminal GB1-tag | (Zhou and Wagner 2010) |
| pMstX | Protein overexpression with N-terminal mistic tag | (This study) |
| pMS73 | CRISPR/Cas9 vector for mutliplexed genome editing in *Ustilago maydis*. Addgene #110629 | (Schuster et al. 2018) |
| p123 | Chromosomal integration plasmid for *U. maydis* | (Aichinger et al. 2003) |
| pFA001 | pMS73 with sgRNA to disrupt *UMAG_04185* | (This study) |
| pFA003 | pMS73 with sgRNA to disrupt *UMAG_01689* | (This study) |
| pFA004 | pMS73 with sgRNA to disrupt *UMAG_10491* | (This study) |
| pFA005 | pMS73 with sgRNA to disrupt *UMAG_01713* | (This study) |
| pFA006 | pMS73 with sgRNA to disrupt *UMAG_00032* | (This study) |
| pFA007 | pMS73 with sgRNA to disrupt *UMAG_03474* | (This study) |
| pFA508 | pEMGB1-*vmp2_118-337_* | (This study) |
| pFA538 | pMS73 with sgRNA to disrupt *UMAG_01689* for deletion of the CTD | (This study) |
| pFA511 | p123-P_00032_-*UMAG_00032* | (This study) |
| pFA512 | p123-P_01689_-*UMAG_01689* | (This study) |
| pFA625 | pMS73 with sgRNA to disrupt *UMAG_01689* for deletion of the NTD | (This study) |
| pFA659 | pMstX-*vmp2* | (This study) |
| pFA670 | pMstX-*vmp1* | (This study) |

**Table S2.** Oligonucleotides used in this study.

| **Target** | **Forward** | **Reverse** |
| --- | --- | --- |
| pFA001 | CAAAATTCCATTCTACAACGGCACGAGCAGCAGCAGTAGAGTTTTAGAGC | CGGCGTTCGACTCTT |
| pFA004 | CAAAATTCCATTCTACAACGGCTCGGAAATCCACACGTTTGTTTTAGAGC | CGGCGTTCGACTCTT |
| pFA005 | CAAAATTCCATTCTACAACGGGCAGAGAGAATGTGCGAGAGTTTTAGAGC | CGGCGTTCGACTCTT |
| pFA007 | CAAAATTCCATTCTACAACGGCCTTTGTCACCAACTTTGCGTTTTAGAGC | CGGCGTTCGACTCTT |
| pFA006 | CAAAATTCCATTCTACAACGGTCTTAATATGCCGACAGATGTTTTAGAGC | CGGCGTTCGACTCTT |
| pFA003 | CAAAATTCCATTCTACAACGGCTGTTCAGGGAGCTGCCGCGTTTTAGAGC | CGGCGTTCGACTCTT |
| pFA508 | AGGAGGGTCTCCCATGGGCAACTACAACCTCGGTAAA | AGGAGGGTCTCCTCGAGTTAGCCGCTGGCGTCGCCAGCGTCGCCAGCATC |
| pFA511 | AGGAGGGTACCGATACAGGACGTTGCTTTTG | AGGAGGCGGCCGCTCAGTACATCTCAGCGCCAT |
| pFA512 | AGGAGGGTACCTCGTATATCAATATATTCTTTGCAT | AGGAGGCGGCCGCTCAGCCGCTGGCGTCGCCAGCGTCGCCAGCATC |
| pFA538 | CAAAATTCCATTCTACAAC**G**GGTAACGACGAAGGCGATGCGTTTTAGAGC | CGGCGTTCGACTCTT |
| pFA625 | CAAAATTCCATTCTACAACGGGAGAGCGAGGTGTCGCGCAGTTTTAGAGC | CGGCGTTCGACTCTT |
| pFA670 | AGGAGGGTCTCCCATGGGCCAGCCGGTTCTGCACATC | AGGAGGGTCTCCTCGAGTTAGTACATTTCAGCACCGTTGG |
| pFA659 | AGGAGGGTCTCCCATGGGCGCGGAATCGCAGTGTGAAGA | AGGAGGGTCTCCTCGAGTTAGCCGCTGGCGTCGCCAGCGTCGCCAGCATC |

**Table S3.** Strains used in this study.

| **Strain** | **Parental strain** | **Vector** | **Genotype** | **Citation** |
| --- | --- | --- | --- | --- |
| SG200 | - | - | - | (Kämper et al. 2006) |
| FA001 | SG200 | pFA001 | ΔUMAG_04185 | This study |
| FA006 | SG200 | pFA003 | ΔUMAG_01689 | This study |
| FA008 | SG200 | pFA004 | ΔUMAG_10491 | This study |
| FA012 | SG200 | pFA005 | ΔUMAG_01713 | This study |
| FA017 | SG200 | pFA006 | ΔUMAG_00032 | This study |
| FA031 | SG200 | pFA007 | ΔUMAG_03474 | This study |
| FA068 | FA006 | pFA512 | ΔUMAG_01689  p123-P_UMAG_01689_-*UMAG_01689* | This study |
| FA071 | FA017 | pFA511 | ΔUMAG_00032  p123-P_UMAG_00032_-*UMAG_00032* | This study |
| FA096 | FA006 | pFA538 | UMAG_01689ΔCTD | This study |
| FA100 | FA006 | pFA625 | UMAG_01689ΔNTD | This study |

**Table S4.** Donor DNA used for CRISPR-Cas9 knockouts in this study.

| **Target** | **Sequence** |
| --- | --- |
| ΔUMAG_04185 | TTTGCACCTTTCCTCGTAAATCGTTTCGAGCTCTCTCACATCGGAGTTGCAGATTAGGTCCTTGTTCGTCAGTTCTGTTA |
| ΔUMAG_10491 | TGGTTTCTCTGTCCTCAATCCTCGTTCGACTTTCATCACACGCGTTCATCTTCGTCCTCCTACTTTGCCATTCGTGATTT |
| ΔUMAG_01713 | CAAGCCAACGCAGAGAGGCCTGAGCGAGAGTCATTGTTCCATTGCTGAGGAGGCATTCACGATTGTCAGCGTTTCTCTCC |
| ΔUMAG_03474 | GCTAGACGCGTATTTGTGCTTGGCGCTGCAGCAGCCAATCCCTCGCTCTTACATACCACGCACATGCGCGTTTGTATTTC |
| Δvmp1 | GTTATCGTCTTGATCTGACCAGCTCGAGCGCGCGACTACGAGCTAGCTGCTCGCTCAGCACATGCCTCATAGGGATGTCA |
| Δvmp2 | CGCTTCAGAGGCTGAACTCGCATGGAACATGGTGGTCAGCGTCTATTCTATTTCAACAATCATTTTTGTTCTGCATGCTG |
| vmp2ΔNTD | GCTGTTGATGATTATTTGGTGGAGCATCGTCTCTGCAGCGGACATCGAATGGATGGGCTTCTACGGCTCCCTCCTCGCTG |
| vmp2ΔCTD | GCTCTCCATCCTCGGAATCATCACCAACTACAACCTCGGTCTGAGTCTATTCTATTTCAACAATCATTTTTGTTCTGCATG |

**Table S5.** Raw data for the statistical analysis of plant infection assays.

| RAW Data | | | | | | | | | | | | | | | |
| --- | --- | --- | --- | --- | --- | --- | --- | --- | --- | --- | --- | --- | --- | --- | --- |
|  | category 0 | | category 1 | | category 2 | | category 3 | | category 4 | | category 5 | | category 6 | |  |
|  | no symptoms | | chlorosis | | ligula swelling | | small tumors | | normal tumors | | heavy tumors | | dead plants | | total plants |
| SG200 | 3,00 | | 3,00 | | 0,00 | | 45,00 | | 39,00 | | 44,00 | | 2,00 | | 136 |
| UMAG_04185 | 2,00 | | 12,00 | | 0,00 | | 29,00 | | 49,00 | | 48,00 | | 4,00 | | 144 |
| UMAG_10491 | 2,00 | | 13,00 | | 0,00 | | 48,00 | | 39,00 | | 37,00 | | 7,00 | | 146 |
| UMAG_01713 | 2,00 | | 6,00 | | 0,00 | | 23,00 | | 47,00 | | 59,00 | | 6,00 | | 143 |
| UMAG_03474 | 1,00 | | 9,00 | | 0,00 | | 40,00 | | 36,00 | | 48,00 | | 8,00 | | 142 |
| UMAG_00032 | 60,00 | | 72,00 | | 0,00 | | 0,00 | | 0,00 | | 0,00 | | 0,00 | | 132 |
| UMAG_01689 | 8,00 | | 78,00 | | 15,00 | | 47,00 | | 0,00 | | 0,00 | | 0,00 | | 148 |
| RAW Data (Percentage) | | | | | | | | | | | | | | | |
|  | category 0 | | category 1 | | category 2 | | category 3 | | category 4 | | category 5 | | category 6 | |  |
|  | no symptoms | | chlorosis | | ligula swelling | | small tumors | | normal tumors | | heavy tumors | | dead plants | |  |
| SG200 | 0,022 | | 0,022 | | 0,000 | | 0,331 | | 0,287 | | 0,324 | | 0,015 | |  |
| UMAG_04185 | 0,014 | | 0,083 | | 0,000 | | 0,201 | | 0,340 | | 0,333 | | 0,028 | |  |
| UMAG_10491 | 0,014 | | 0,089 | | 0,000 | | 0,329 | | 0,267 | | 0,253 | | 0,048 | |  |
| UMAG_01713 | 0,014 | | 0,042 | | 0,000 | | 0,161 | | 0,329 | | 0,413 | | 0,042 | |  |
| UMAG_03474 | 0,007 | | 0,063 | | 0,000 | | 0,282 | | 0,254 | | 0,338 | | 0,056 | |  |
| UMAG_01689 | 0,054 | | 0,527 | | 0,101 | | 0,318 | | 0,000 | | 0,000 | | 0,000 | |  |
| UMAG_00032 | 0,455 | | 0,545 | | 0,000 | | 0,000 | | 0,000 | | 0,000 | | 0,000 | |  |
| Standard devaiation (Percentage) | | | | | | | | | | | | | | | |
|  | category 0 | | category 1 | | category 2 | | category 3 | | category 4 | | category 5 | | category 6 | |  |
|  | no symptoms | | chlorosis | | ligula swelling | | small tumors | | normal tumors | | heavy tumors | | dead plants | |  |
| SG200 | 0,021 | | 0,022 | | 0,000 | | 0,083 | | 0,018 | | 0,120 | | 0,012 | |  |
| UMAG_04185 | 0,012 | | 0,021 | | 0,000 | | 0,024 | | 0,073 | | 0,021 | | 0,048 | |  |
| UMAG_10491 | 0,023 | | 0,085 | | 0,000 | | 0,070 | | 0,059 | | 0,048 | | 0,069 | |  |
| UMAG_01713 | 0,012 | | 0,058 | | 0,000 | | 0,086 | | 0,078 | | 0,072 | | 0,036 | |  |
| UMAG_03474 | 0,012 | | 0,002 | | 0,000 | | 0,028 | | 0,022 | | 0,048 | | 0,026 | |  |
| UMAG_00032 | 0,109 | | 0,109 | | 0,000 | | 0,000 | | 0,000 | | 0,000 | | 0,000 | |  |
| UMAG_01689 | 0,032 | | 0,139 | | 0,109 | | 0,271 | | 0,000 | | 0,000 | | 0,000 | |  |
| Student’s t-test | | | | | | | | | | | | | | | |
|  | | category 0 | | category 1 | | category 2 | | category 3 | | category 4 | | category 5 | | category 6 | |
|  | | no symptoms | | chlorosis | | ligula swelling | | small tumors | | normal tumors | | heavy tumors | | dead plants | |
| SG200 vs 00032 | | 0,003 | | 0,002 | | - | | 0,004 | | 0,000 | | 0,005 | | 0,116 | |
| SG200 vs 01689 | | 0,189 | | 0,002 | | 0,171 | | 0,939 | | 0,000 | | 0,005 | | 0,116 | |

| RAW Data Vmp1 | | | | | | | | |
| --- | --- | --- | --- | --- | --- | --- | --- | --- |
|  | category 0 | category 1 | category 2 | category 3 | category 4 | category 5 | category 6 |  |
|  | no symptoms | chlorosis | ligula swelling | small tumors | normal tumors | heavy tumors | dead plants | total plants |
| SG200 | 0,00 | 6,00 | 0,00 | 63,00 | 78,00 | 3,00 | 0,00 | 150,00 |
| ∆Vmp1 | 60 | 72 | 0 | 0 | 0 | 0 | 0 | 132,00 |
| compl. Vmp1 | 0,00 | 16,00 | 0,00 | 123,00 | 1,00 | 2,00 | 1,00 | 143,00 |
| RAW Data (Percentage) | | | | | | | | |
|  | category 0 | category 1 | category 2 | category 3 | category 4 | category 5 | category 6 |  |
|  | no symptoms | chlorosis | ligula swelling | small tumors | normal tumors | heavy tumors | dead plants |  |
| SG200 | 0,000 | 0,040 | 0,000 | 0,420 | 0,520 | 0,020 | 0,000 |  |
| ∆Vmp1 | 0,455 | 0,545 | 0,000 | 0,000 | 0,000 | 0,000 | 0,000 |  |
| compl. Vmp1 | 0,000 | 0,112 | 0,000 | 0,860 | 0,007 | 0,014 | 0,007 |  |

**Table S5 (continued).** Raw data for the statistical analysis of plant infection assays.

| RAW Data Vmp2 | | | | | | | | |
| --- | --- | --- | --- | --- | --- | --- | --- | --- |
|  | category 0 | category 1 | category 2 | category 3 | category 4 | category 5 | category 6 |  |
|  | no symptoms | chlorosis | ligula swelling | small tumors | normal tumors | heavy tumors | dead plants | total plants |
| SG200 | 0,00 | 5,00 | 0,00 | 25,00 | 39,00 | 60,00 | 5,00 | 134,00 |
| ∆Vmp2 | 8 | 78 | 15 | 47 | 0 | 0 | 0 | 148,00 |
| compl. Vmp2 | 0,00 | 0,00 | 1,00 | 50,00 | 72,00 | 18,00 | 3,00 | 144,00 |
| ∆NTD | 4,00 | 29,00 | 6,00 | 24,00 | 2,00 | 0,00 | 0,00 | 65,00 |
| ∆CTD | 6,00 | 42,00 | 4,00 | 9,00 | 0,00 | 0,00 | 0,00 | 61,00 |
| RAW Data (Percentage) | | | | | | | | |
|  | category 0 | category 1 | category 2 | category 3 | category 4 | category 5 | category 6 |  |
|  | no symptoms | chlorosis | ligula swelling | small tumors | normal tumors | heavy tumors | dead plants |  |
| SG200 | 0,000 | 0,037 | 0,000 | 0,187 | 0,291 | 0,448 | 0,037 |  |
| ∆Vmp2 | 0,054 | 0,527 | 0,101 | 0,318 | 0,000 | 0,000 | 0,000 |  |
| compl. Vmp2 | 0,000 | 0,000 | 0,007 | 0,347 | 0,500 | 0,125 | 0,021 |  |
| ∆NTD | 0,062 | 0,446 | 0,092 | 0,369 | 0,031 | 0,000 | 0,000 |  |
| ∆CTD | 0,098 | 0,689 | 0,066 | 0,148 | 0,000 | 0,000 | 0,000 |  |

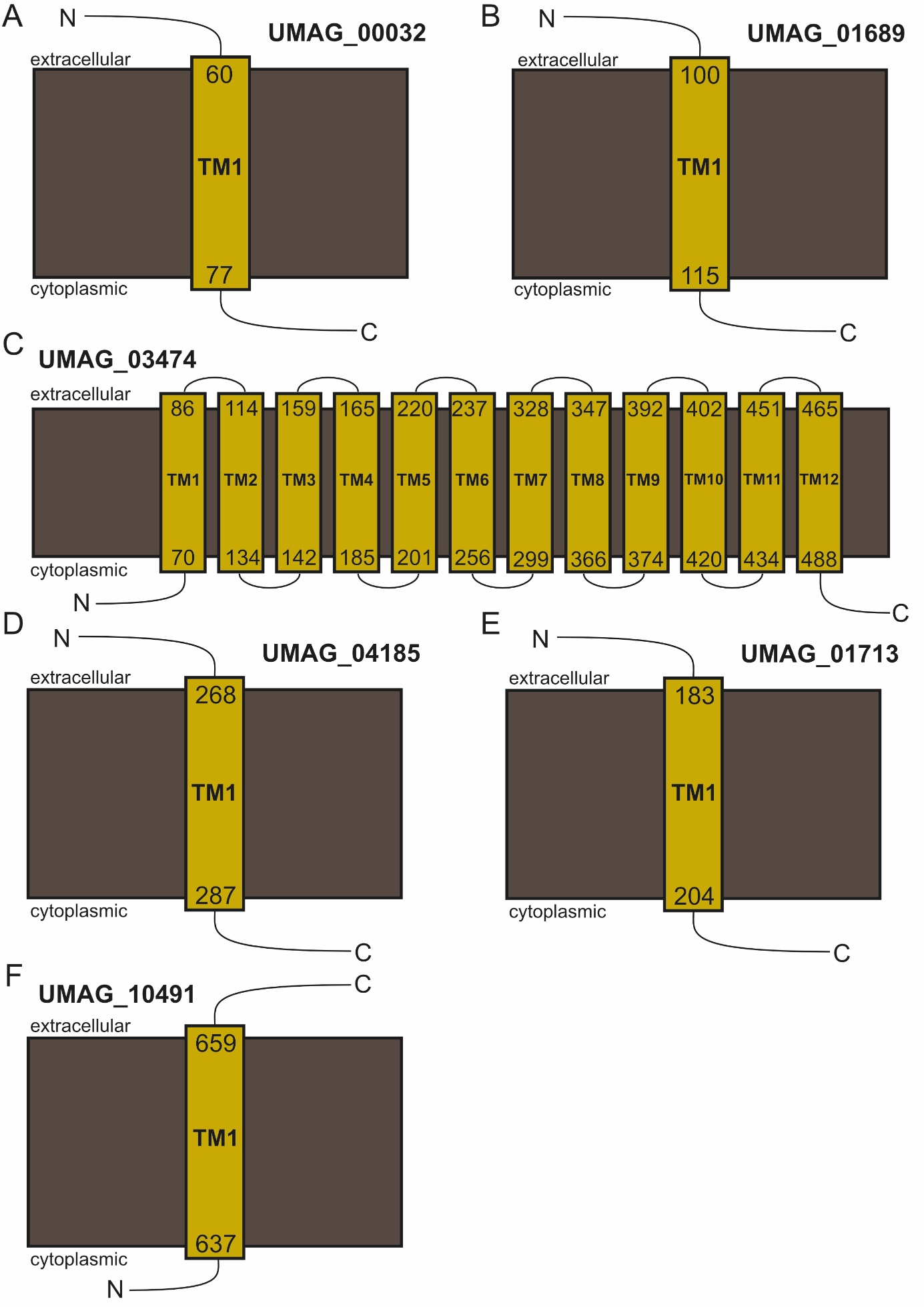

**Figure S1.** **Prediction of transmembrane helices.** Schematic representation of the residues predicted to be involved in the formation of transmembrane helices (TM) of all six proteins (**A**-**F**) according to the Consensus Constrained TOPology prediction web server CCTOP and PSIPRED (Dobson, Reményi, and Tusnády 2015; Buchan and Jones 2019). Cytoplasmic and extracellular regions are labeled accordingly whereas N- and C-terminal regions are labeled with N and C, respectively.

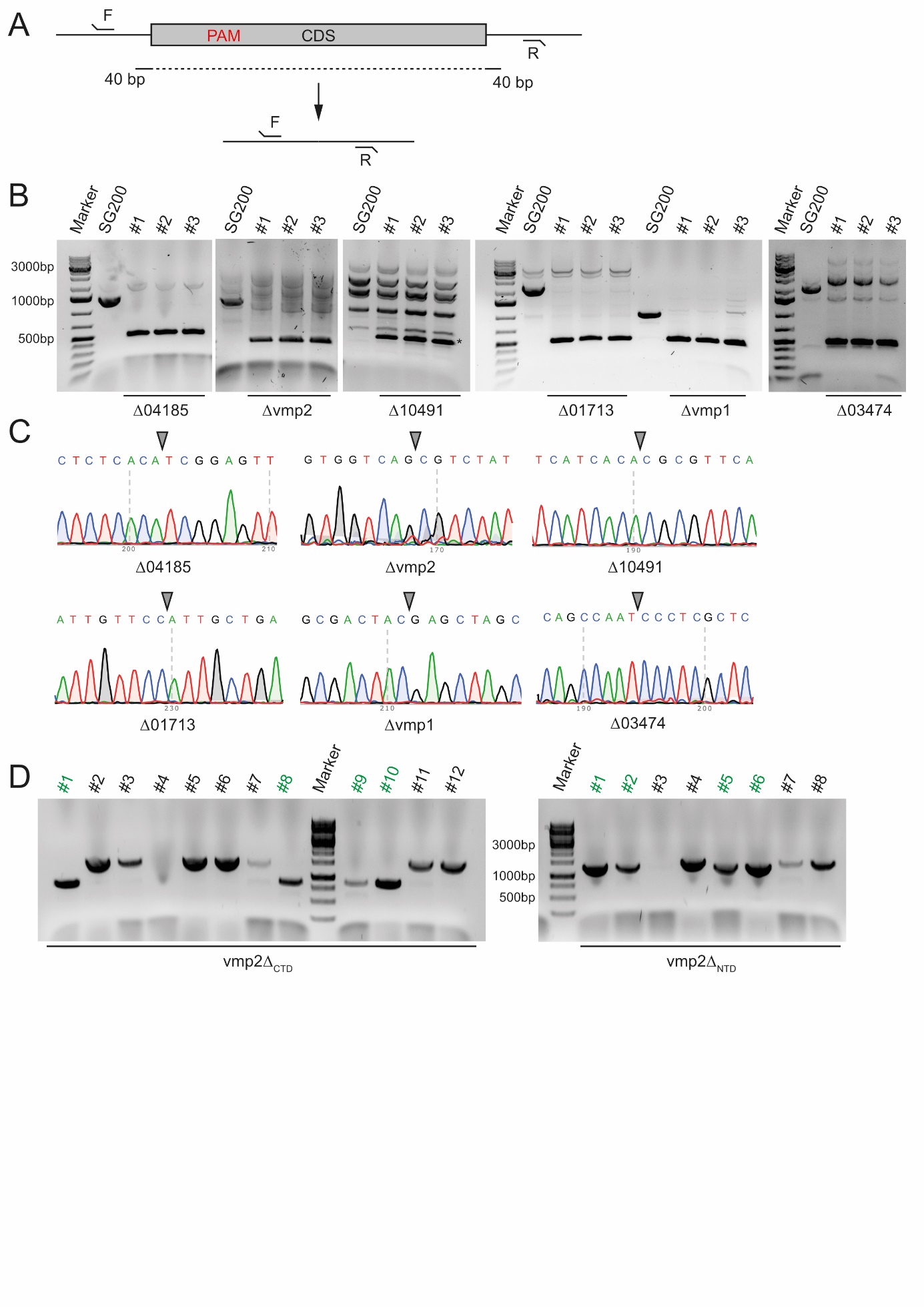

**Figure S2.** **Evaluation of the CRISPR-Cas9-based knockout approach.** **A.** Knockout approach using a donor DNA of 80 bp leading to a clean deletion of the respective open reading frames (ORFs). **B.** Colony-PCR of three individual transformants of each deletion. In case of a successful replacement of the respective gene with the donor DNA, a PCR product of 500 bp can be observed. **C.** Sequencing results of one transformant for each strain showing the clean deletion of the respective ORFs. The triangle indicates the former ORF. **D.** Colony PCR of transformants deleted for the region encoding the CTD and the NTD of Vmp2, respectively. Positive transformants are colored in green. The PCR products were sequenced to evaluate that an intact ORF is present in all transformants.

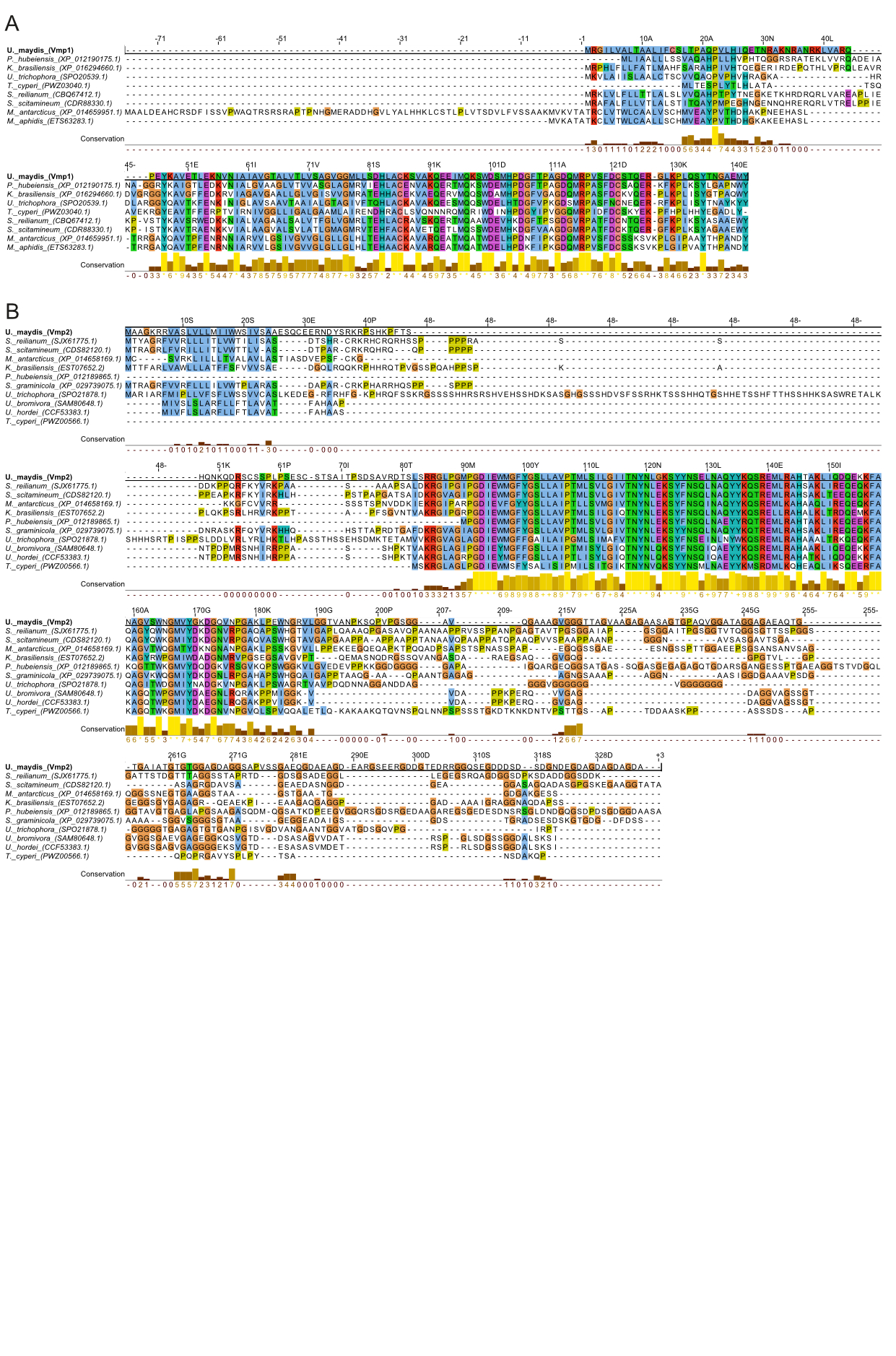

**Figure S3.** **Sequence alignment of Vmp1 and Vmp2 to homologs from related smut fungi**. Homologous proteins of both Vmp1 (**A**) and Vmp2 (**B**) were identified using the BlastP (Basic Local Alignment Search Tool) webserver and the obtained sequences were aligned using the Clustal Omega webserver (Sievers et al. 2011). Visualization and conservation determination was done with Jalview. For the coloring of the individual residues the default scheme from Clustal Omega was used. The conservation score was determined by measuring the number of conserved physico-chemical properties for each column of the alignment. Its calculation is based on the one used in the AMAS (Analysis of Multiply Aligned Sequences) method of multiple sequence alignment analysis (Livingstone and Barton 1993).

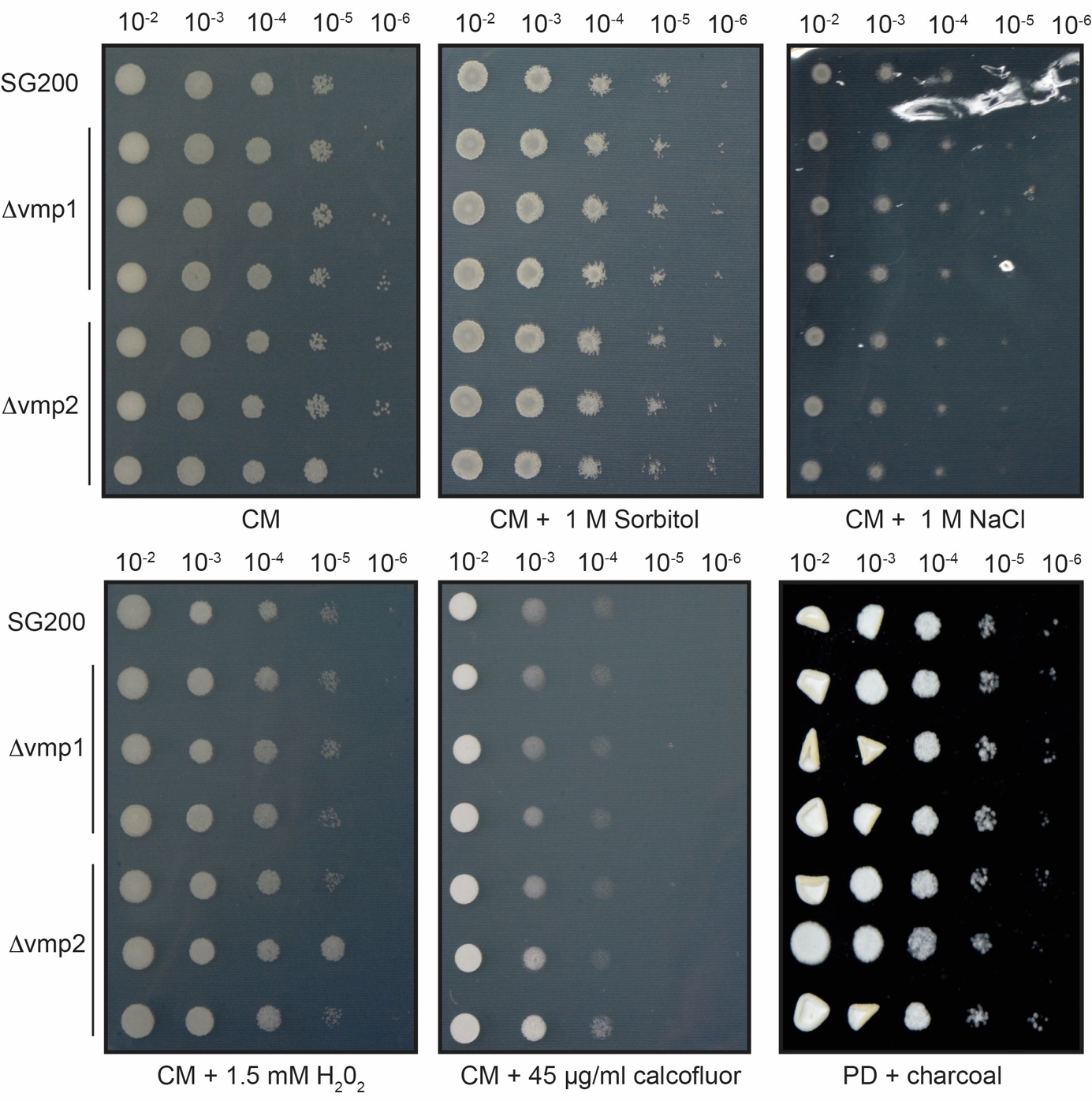

**Figure S4. Stress assays of the vmp1 and vmp2 deletion strains.** Multiple stress assays and for filamentation of SG200, SG200Δvmp1 and SG200Δvmp2 on CM plates containing Sorbitol, NaCl, H2O2 and Calcofluor white and potato dextrose (PD) plates containing charcoal.

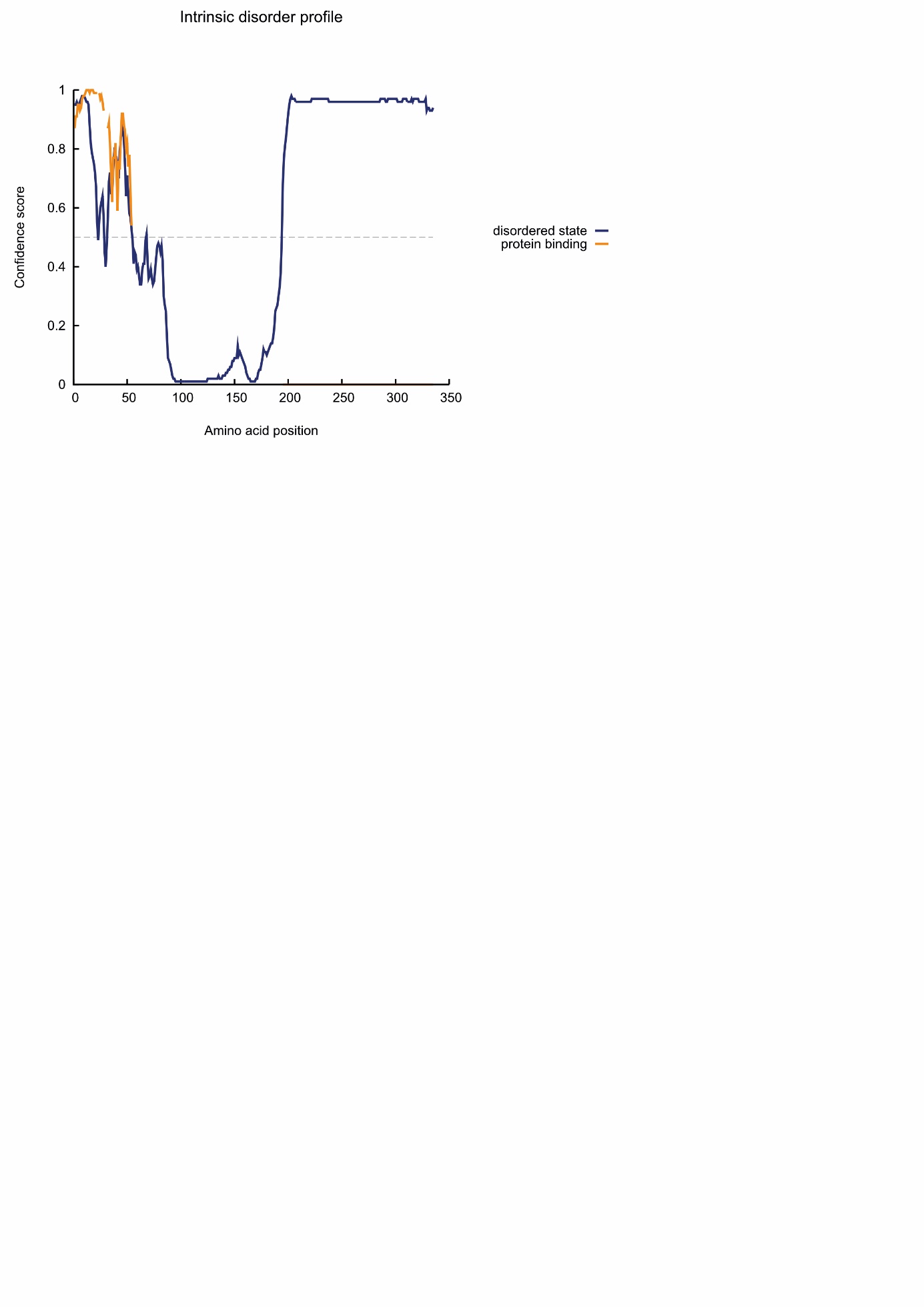

**Figure S5**. **Intrinsic disorder profile of Vmp2.** Diagram was obtained from the PSIPRED web server (Buchan and Jones 2019).
